# Planning Face, Hand, and Leg Movements: Anatomical Constraints on Preparatory Inhibition

**DOI:** 10.1101/529107

**Authors:** Ludovica Labruna, Claudia Tischler, Cristian Cazares, Ian Greenhouse, Julie Duque, Florent Lebon, Richard B. Ivry

**Author notes:** **Corresponding author:** Ludovica Labruna, Department of Psychology, 2121 Berkeley Way, University of California, Berkeley, CA 94720.

## Abstract

Motor-evoked potentials (MEPs), elicited by Transcranial Magnetic Stimulation (TMS) over the motor cortex, are reduced during the preparatory period in delayed response tasks. Here we examine how MEP suppression varies as a function of the anatomical organization of the motor cortex. MEPs were recorded from a left index muscle while participants prepared a hand or leg movement in Experiment 1, or prepared an eye or mouth movement in Experiment 2. In this manner, we assessed if the level of MEP suppression in a hand muscle varied as a function of the anatomical distance between the agonist for the forthcoming movement and the muscle targeted by TMS. MEPs suppression was attenuated when the cued effector was anatomically distant from the hand (e.g., leg or facial movement compared to finger movement). A similar effect was observed in Experiment 3 in which MEPs were recorded from a muscle in the leg and the forthcoming movement involved the upper limb or face. These results demonstrate an important constraint on preparatory inhibition: It is sufficiently broad to be manifest in a muscle that is not involved in the task, but is not global, showing a marked attenuation when the agonist muscle belongs to a different segment of the body.

**New & Noteworthy:** Using TMS, we examine changes in corticospinal excitability as people prepare to move. Consistent with previous work, we observe a reduction in excitability during the preparatory period, an effect observed in both task relevant and task irrelevant muscles. However, this preparatory inhibition is anatomically constrained, attenuated in muscles belonging to a different body segment than the agonist of the forthcoming movement.

## Introduction

Transcranial Magnetic Stimulation (TMS) has proven to be a powerful tool to assess the dynamics of corticospinal (CS) excitability during response preparation in humans (Bestmann and Duque 2016; Cos et al. 2014; Klein et al. 2012; Leocani et al. 2000). In a delayed response task, a cue is provided to indicate the forthcoming response, with that response initiated after the onset of an imperative signal (e.g., cue left or right index finger movement). TMS studies have shown local increases in cortical excitability in primary motor cortex (M1) during the delay period (Davranche et al. 2007; Duque and Ivry 2009; Tandonnet et al. 2010), as well as broad suppression of cortico-spinal excitability during the same time window (Greenhouse et al. 2015b; Hannah et al. 2018). Indeed, when single-pulse TMS is delivered over the primary motor cortex during the delay period, motor evoked potentials (MEPs) elicited from the targeted muscle show profound suppression, regardless of whether that muscle is required to perform the cued movement (i.e., selected) or not required for the forthcoming response (non-selected) (Duque et al. 2017; Duque and Ivry 2009; Quoilin et al. 2016). After the imperative, the selected muscle shows a rapid increase in excitability, while the non-selected muscles remain suppressed (Duque et al. 2014; Duque and Ivry 2009; Klein et al. 2016).

Interestingly, in many studies, the strongest level of MEP suppression during the delay period is observed when the muscle is the agonist for the selected response (Duque and Ivry 2009; Klein et al. 2016). This observation led to the hypothesis that preparatory inhibition is designed to prevent premature movement. In contrast, MEP suppression of non-selected muscles has been considered a useful mechanism for action selection, helping to sharpen the preparation of a selected movement by inhibiting alternative representations (Duque et al. 2005, 2010; Leocani et al. 2000; Tandonnet et al. 2011). This process might be implemented by inhibitory interactions between competing alternatives or might rely on a more generic form of inhibition whereby the choice of one action is accompanied by broad inhibition of the motor system to lower interference from irrelevant motor representations (Duque et al. 2017).

Using a reaction time (RT) task in which the response was fixed for an entire block of trials, Greenhouse et al. (2015b) observed substantial preparatory inhibition in task-irrelevant muscles (e.g., in a left index finger agonist when the right pinky was always used to make the response). Indeed, the magnitude of the MEP suppression was similar in task-irrelevant muscles compared to task-relevant muscles (e.g., in a left index finger agonist when the cued response was either the left index finger or the left pinky). These findings are difficult to reconcile with the hypothesis that preparatory inhibition assists action selection, and points to a more generic process.

However, other findings suggest that preparatory inhibition is not generic. In choice RT tasks, the magnitude of MEP suppression in a non-selected muscle varies as a function of the relationship between the members of the response set: A bigger reduction in excitability is found when the response set involves homologous effectors compared to when the response set involves non-homologous effectors (Duque et al. 2014; Labruna et al. 2014), a result that may reflect functional or anatomical links between homologous representations across the two hemispheres (van den Heuvel and Hulshoff Pol 2010). Similarly, MEP suppression in a non-selected muscle (e.g., left index finger) is greater when the planned movement involves another upper limb effector (e.g., right index finger) compared to when the planned movement involves a lower limb effector (e.g. right leg).

Taken together, these findings suggest that preparatory inhibition is subject to anatomical constraints. To further explore this hypothesis, we systematically manipulated the response set to derive comparisons between conditions in which the probed muscle was close or distant to the members of the response set in terms of anatomy or function. An overview of the experimental plan is presented in Fig 1. In Exps 1 and 2, MEPs were always elicited from the left first dorsal interosseous (FDI) muscle, the agonist for left index finger abduction movements. We created conditions in which this muscle was selected or not selected for the forthcoming response or task-irrelevant. Of primary interest, we manipulated the response set to examine whether corticospinal excitability in left FDI varied as a function of the other candidate movements, choosing a range of movements that involved the same or different side of the body or same or different body segment.

**Figure 1.**
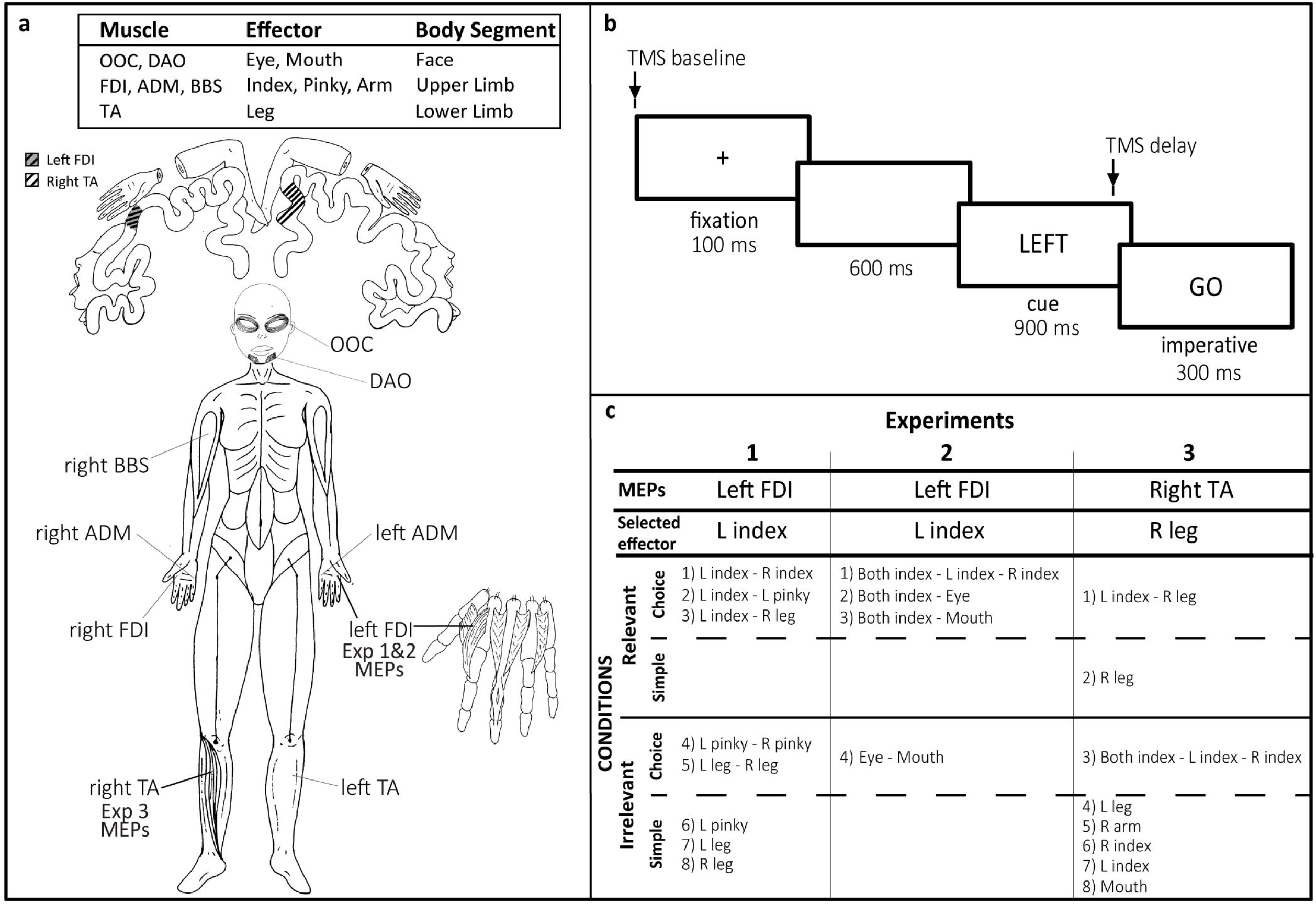
Overview of the three experiments. (a) Primary agonist muscles used for the responses. A schematic of the cortical homunculus is shown on top, with highlighted regions (diagonal lines) indicating approximate location of the left hand FDI, the muscle targeted for TMS in Exps 1 and 2, and right TA, the muscle targeted for TMS in Exp 3. Abbreviations: orbicularis oculi (OOC), depressor anguli oris (DAO), first dorsal interossus (FDI), abductor digiti minimi (ADM), biceps brachii short (BBS), tibialis anterior (TA). (b) Sequence of events in the delayed response task. The TMS pulse was either coincident with the onset of the fixation cross (TMS baseline) or occurred 800 ms into the delay period (TMS delay). (c) Response set for each condition in the experiments. Relevant and Irrelevant refer to conditions in which the targeted muscle was either part of, or not part of the response set. In Simple conditions, the same movement was cued on each trial, whereas in Choice conditions, the cue specified the forthcoming movement.

In Exp 1, this involved a comparison of left FDI MEPs between different sets of hand and leg movements. Here we sought to replicate our earlier findings (Duque et al. 2014; Labruna et al. 2014) showing that reduced excitability is modulated by anatomical similarity, but not by task relevance. In particular, we expected to observe greater MEP suppression in the left FDI when the selected response involved a finger movement not requiring left FDI, compared to when the selected response involved a leg movement. In Exp 2, the focus was on response sets in which index finger movements were paired with either eye or mouth movements. By combining hand and facial movements, we obtain a second test of inter-segmental interactions in preparatory inhibition. To ensure that our results are not specific to hand muscles, the TMS probe was targeted at a lower leg muscle, the right tibialis anterior (TA) muscle, in Exp 3. The focus here was to determine if the patterns of intra- and intersegmental interactions observed in a hand muscle would be similar in a leg muscle.

## Methods

### Participants

Thirty-six healthy, right-handed participants (Oldfield, 1971) were tested, 12 in each experiment (mean ± SD: Exp 1: 20.8 ± 1.1 years old, 10 men; Exp 2: 22.6 ± 5.4 years old, 6 men; Exp 3: 21.0 ± 1.7 years old, 5 men). Participants were recruited from a website maintained by UC Berkeley to assist investigators in identifying individuals willing to participate in scientific research. The participants were naïve to the purpose of the study and were financially compensated for their participation. The recruitment process used in the present study excluded professional musicians or individuals with an extensive history of experience in playing a musical instrument. The protocol was approved by the institutional review board of the University of California, Berkeley. As part of the informed consent, participants completed a TMS safety checklist prior to the start of the experiment.

## Procedure

### TMS

In all experiments, participants sat in front of a computer screen with both hands resting on a pillow, palms down, with the arms relaxed in a semi-flexed position. TMS was applied over the M1 during a delayed response task to measure changes in the excitability state of the CS pathway during response preparation. The TMS was positioned to elicit MEPs in a single targeted muscle across all conditions in a given experiment (for a review of general procedures used to measure corticospinal excitability during response preparation, see Bestmann and Duque 2016; Duque et al. 2017).

TMS pulses were delivered with a monophasic Magstim 200^2^ magnetic stimulator (Magstim, Whitland, Dyfed, UK). In Exps 1 and 2, a 90 mm figure-of-eight coil was positioned over the participant’s scalp above the right M1. The coil was placed tangentially, in the posterior-anterior direction, with the handle oriented toward the back of the head, and laterally at a 45° angle from the midline, an orientation that is approximately perpendicular to the central sulcus. We identified the optimal position to elicit MEPs in the left FDI muscle. In Exp 3, the coil was positioned to optimize MEPs in the TA of the right leg, the agonist for adduction movements of the right foot. Given that the leg region is in the depth of the sulcal, we used a 110 mm double cone coil that produces a higher induced current (Deng et al. 2014). The coil was positioned over the left M1, in a posterior–anterior orientation, 1 cm above and 1 cm to the right of the vertex.

Once identified, the optimal position for eliciting MEPs in the targeted muscle (left FDI or right TA) was marked on the scalp to provide a reference point for the experimental session. The participant’s resting motor threshold (rMT) was identified at the hotspot and defined as the minimum TMS intensity required to evoke MEPs of ~50 μV peak-to-peak amplitude on 5 of 10 consecutive trials (Rossini et al., 1994). Averaging across Exps 1 and 2, the mean rMT for the left FDI corresponded to 45% (SD = 7) of maximum stimulator output (MSO). In Exp 3, the mean rMT for the right TA was 78% (SD = 13) of MSO. The intensity of TMS was set to 115% of the individual rMT.

### EMG Recording

EMG was recorded with surface electrodes placed above selected muscles (see below). The EMG signal was continuously monitored on-line to ensure that participants maintained a relaxed posture over the course of the experiment. The EMG signals were amplified and bandpass-filtered on-line between 20 and 450 Hz (Delsys, Inc.). The signals were digitized at 2000 Hz for off-line analysis.

In Exp 1, six EMG electrodes were used, positioned to record from FDI, abductor digiti minimi (ADM) and TA on both sides. In Exp 2, we used four electrodes. Two were placed on the left and right FDI. The other two were placed on the face, one over the left orbicularis oculi (OOc) and the other over the left depressor anguli oris (DAO), to record EMG for eye and mouth muscles, respectively. We only considered activity on one side given that movements with the face effectors, when produced, entailed a relatively symmetric activation in the left and right side muscles (Cattaneo and Pavesi 2014). In Exp 3, six electrodes were used to record activity from FDI and TA bilaterally, and from left DAO (mouth muscle) and the short head of right biceps brachii (BBS), the agonist for arm flexion.

### Delayed-response task

A delayed response task was used to study changes in corticospinal excitability during response preparation (Fig 1b). Each trial began with the brief presentation (100 ms) of a cross at the center of the computer monitor, followed by a 600 ms blank screen and then the presentation of a preparatory cue for 900 ms. The cue consisted of one or two words, positioned at the screen center, specifying the effector for the forthcoming response (e.g., “LEFT”, see below). At the end of the 900 ms delay period, the word “GO” appeared for 300 ms, providing a signal to the participant to produce the cued response. The participants were instructed to prepare their response during the delay period in order to respond as quickly as possible once the imperative stimulus appeared.

A single TMS pulse was applied on each trial. The pulse was either coincident with the onset of the fixation cross (TMS baseline) or occurred 100 ms before the imperative, 800 ms into the delay period (TMS delay). The TMS baseline and delay trials were randomized, with the constraint that the two timings occurred equally often for each cue. The variation in MEP amplitudes at TMS delay with respect to TMS baseline provided a probe of changes in CS excitability during movement preparation. Although preparatory time may vary for different movements, the long delay period used here ensures that participants have sufficient time to reach an optimal state of preparation prior to the imperative. The duration of the inter-trial interval (ITI) was variable and fluctuated between 3000-3500 ms. We note that, with this design, the participants can anticipate the TMS pulse during the delay period if it did not occur at baseline. However, prior work in our lab showed that changes in CS excitability during the delay period are not related to the anticipation of a TMS pulse (Greenhouse et al. 2015b).

In each experiment (summarized in Fig 1c), trials were grouped in blocks, with each block involving only one condition. Participants were informed of the response set and their associated cues prior to the start of each block (see below). In Choice RT conditions, there were two or three possible responses and their order was randomized within the block. In Simple RT conditions, the response set consisted of a single response. There were 60 trials in each Choice condition, three of which were catch trials (no imperative). There were 40 trials in each Simple condition, two of which were catch trials. We recorded 20 baseline MEPs for each condition and 20 MEPs for each cue condition in the delay period, a sample size recommended to obtain reliable MEP measures (Biabani et al., 2018). The blocks lasted approximately eight and six minutes for the Choice and Simple RT conditions, respectively. The order of the blocks was randomized across participants (but see constrains in Exp 1).

The muscle from which the MEPs were recorded (left FDI in Exps 1 and 2, right TA in Exp 3) was always relevant or irrelevant for a given block (as highlighted in Fig 1c). The former situation occurred in blocks where the targeted muscle was the agonist for an effector that was part of the response set (and either selected or non-selected on each trial). In contrast, the targeted muscle was irrelevant when it was the agonist for an effector that was not part of the response set in the block. We use the terminology task-relevant and task-irrelevant blocks to describe this aspect of the design.

#### Experiment 1

In Exp 1, we examined CS excitability changes in left FDI as the participants prepared movements with either the left or right hand/leg. There were eight conditions, five of which involved Choice RT tasks. For three of these, left FDI was relevant, with left index finger paired with either the right index finger, the left pinky, or the right leg. These three conditions were selected to compare preparatory inhibition in a hand muscle when the alternative response involved a homologous effector, another effector on the same hand, or an effector of another body segment. For the other two Choice conditions, the left index finger was irrelevant, with the response set consisting of either left/right pinky movements or left/right leg movements. Here we evaluate preparatory inhibition in left FDI when the left index finger is irrelevant but either at the same body segment (intra-segmental) or at a different body segment (inter-segmental) as the effectors included in the response set. Left FDI was also irrelevant in the three Simple RT blocks. These conditions allowed us to ask the same question as with the irrelevant Choice blocks, but without the choice component given that the response was fixed for a given block (left pinky, right or left leg).

When the response set involved a left and right effector, the cues were “Left” and “Right”. When the response set involved two left hand options, the cues were “Index” and “Pinky”. The word “Left” or “Right” was used as the cue in the three Simple RT conditions. Index and pinky responses required an abduction of the specified finger, bringing it away from the center of the hand. For leg responses, the participant produced adduction movements, lifting the foot toward the body midline.

The block order was randomized across participants with the constraint that the left index-right index pairing was always tested last. We did so because we were concerned that some participants might tire over the duration of a 120 min experiment. Given that the left-right index pairing has been used in numerous other studies, we opted to test this one last since the results here could be compared to prior results, providing a crude reliability check.

#### Experiment 2

Exp 2 was designed to further investigate anatomical constraints on preparatory inhibition. A key comparison in Exp 1 involved changes in the MEPs of a hand muscle when preparing a leg movement. In Exp 2, we extended this inter-segmental test, but now examined changes in the MEPs of a hand muscle when preparing a facial movement. Moreover, by comparing different facial gestures, we can assess if the spread of preparatory inhibition is a function of cortical distance. Based on the classic motor homunculus, we would expect MEPs from left FDI would show more suppression when the selected response involves the eye compared to the mouth, given that the eye representation is anatomically closer to the hand area (Fig 1a).

Given that facial movements are generally bilateral (Cattaneo and Pavesi 2014), we thought it important to compare these movements to bilateral hand movements. There were four conditions (Fig 1c), with the order randomized across participants. For three of these, the left FDI was relevant, with bimanual index finger movements combined with either eye or mouth movements, or with unimanual left and right index finger movements. The latter block was used as a control condition to establish a baseline. In the fourth block, the choice was between a mouth and an eye movement, with the left FDI being irrelevant.

Finger movements were cued with the words “Left index”, “Right index”, or “Both index”. Eye and mouth movements were cued with the words “Eyes” or “Mouth”, respectively. Finger responses were as in Exp 1 (index finger abduction). Eye movements consisted of a single volitional squint with both eyes. The mouth movements required the participants to make a volitional smile, with the instruction to show as much of the teeth as possible.

#### Experiment 3

To ensure that the CS excitability changes observed in Exp 1 and 2 were not specific to MEPs elicited in a hand muscle, we targeted the TA muscle of the right leg in Exp 3. MEPs are more difficult to elicit from leg muscles: Not only is the leg region is in the depth of the sulcal, but the motor representations of leg muscles may contain fewer or weaker corticospinal projections (Kesar et al. 2018). Given this challenge, the thresholding phase of Exp 3 also served as a screening procedure: We recruited 23 participants to identify 12 individuals for whom we were able to consistently elicit MEPs in the right TA.

There were a total of eight conditions, with the order randomized across these 12 participants. The right TA muscle was relevant in two conditions, one in which the right leg was tested in a Simple RT task and one in which the right leg was paired with the left index finger in a Choice RT task. Note that we opted to record MEPS from the right TA rather than the left TA given that, by doing so, we have a condition that is identical to one tested in Exp 1 (left index paired with right leg).

The right TA was irrelevant in the other six conditions. Five of these were Simple RT tasks, with the responses made (in separate blocks) with either the mouth, right arm, left index finger, right index finger, or left leg. For the remaining Choice RT condition, we used the 3-choice manual condition of Exp 2 (left, right or bimanual index finger movement).

For the Simple RT blocks, the words “Left Index”, “Right Index”, “Right Arm”, “Mouth” or “Left Leg” were used. In the Choice RT blocks, the cues were “Left Index”, “Right Index”, “Both Index” or “Right Leg”. The required movements for each effector were as in Exps 1 and 2.

### Data and statistical Analysis

The EMG data were analyzed offline using customized routines within Matlab, as well as visual inspection of individual traces to identify artifacts. From the EMG data, we extracted two dependent variables: The peak-to-peak amplitude of the MEP (left FDI in Exps 1 and 2; right TA in Exp 3) and the reaction time (RT). To prevent contamination of the MEP measurements by fluctuations in background EMG, trials were excluded if the background EMG activity was greater than 0.01 mV in the 200-msec window preceding the TMS pulse (Duque et al. 2014; Quoilin et al. 2016; Wilhelm et al. 2016). We also excluded MEPs that were above or below 3 SD of the mean MEP amplitude for that condition, as well as those in which there was EMG activity associated with a non-cued response (selection errors). Overall, 9% of the trials (SD = 2%) were excluded from the analysis (approximately 50% of these were due to the outlier exclusion criterion).

The mean MEP values were calculated for the TMS baseline and delay probes, with the latter calculated separately for each cued effector. To assess CS excitability changes during response preparation, we subtracted the mean delay period MEPs from the mean baseline MEPs on an individual basis and normalized these values by dividing the difference by the mean baseline value. The scores were multiplied by 100 to express as percentage scores, with negative values indicative of preparatory inhibition. Given that many studies have confirmed the existence of preparatory inhibition (for reviews see Bestmann and Duque 2016; Duque et al. 2017), one-tailed t-tests were used in within-condition comparisons to evaluate whether the MEPs were inhibited relative to baseline (i.e., comparison of the normalized scores for each condition to the null hypothesis that the scores would be distributed around zero). The Shapiro-Wilk’s test was used to assess if the scores for a given condition met the normality assumption. When this test indicated a violation of the normality assumption, we analyzed the data with the non-parametric Wilcoxon Signed Rank test.

For comparisons of the preparatory MEP changes between conditions, we used repeated-measures analyses of variance (ANOVA_RM_), with post-hoc tests based on the Bonferroni method, adjusted for multiple comparisons. When the contrast included a condition that violated the assumption of normality, we used the non-parametric Friedman Test, with the Wilcoxon signed-rank test for post-hoc comparisons. The post-hoc tests in Exps 1 and 2 were two-tailed since we did not have strong a priori hypotheses. In Exp 3, a one-tailed test was employed given that the results of the first two experiments led to a test of a specific hypothesis. Effect sizes are reported using partial eta-squared 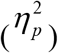 for the ANOVA, and Cohen’s d for the planned contrasts in which the data met the normality assumption. For the non-parametric Wilcoxon signed-rank test, the effect size r was calculated as 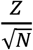 (Rosenthal, 1991). The reported p-values for these are adjusted for multiple comparisons.

RT was defined as the time interval between the onset of the imperative signal and the time point at which the EMG activity of the agonist muscle for the cued response exceeded 3 SD of the mean of the rectified signal for the entire trial epoch. ANOVA_RM_ were also used to analyze these data.

## Results

### CS Excitability

The goal of this study was to explore constraints on preparatory inhibition. We assessed whether changes in corticospinal excitability observed during the delay period varied as a function of the effectors involved in the task and their anatomical relationship with the muscle probed with TMS. To assess whether CS excitability was inhibited during the preparatory period, MEPs elicited during the delay period were compared to MEPs elicited at baseline (i.e., trial onset). A summary of these within-condition comparisons for all three experiments is presented in Table 1.

**Table 1.**
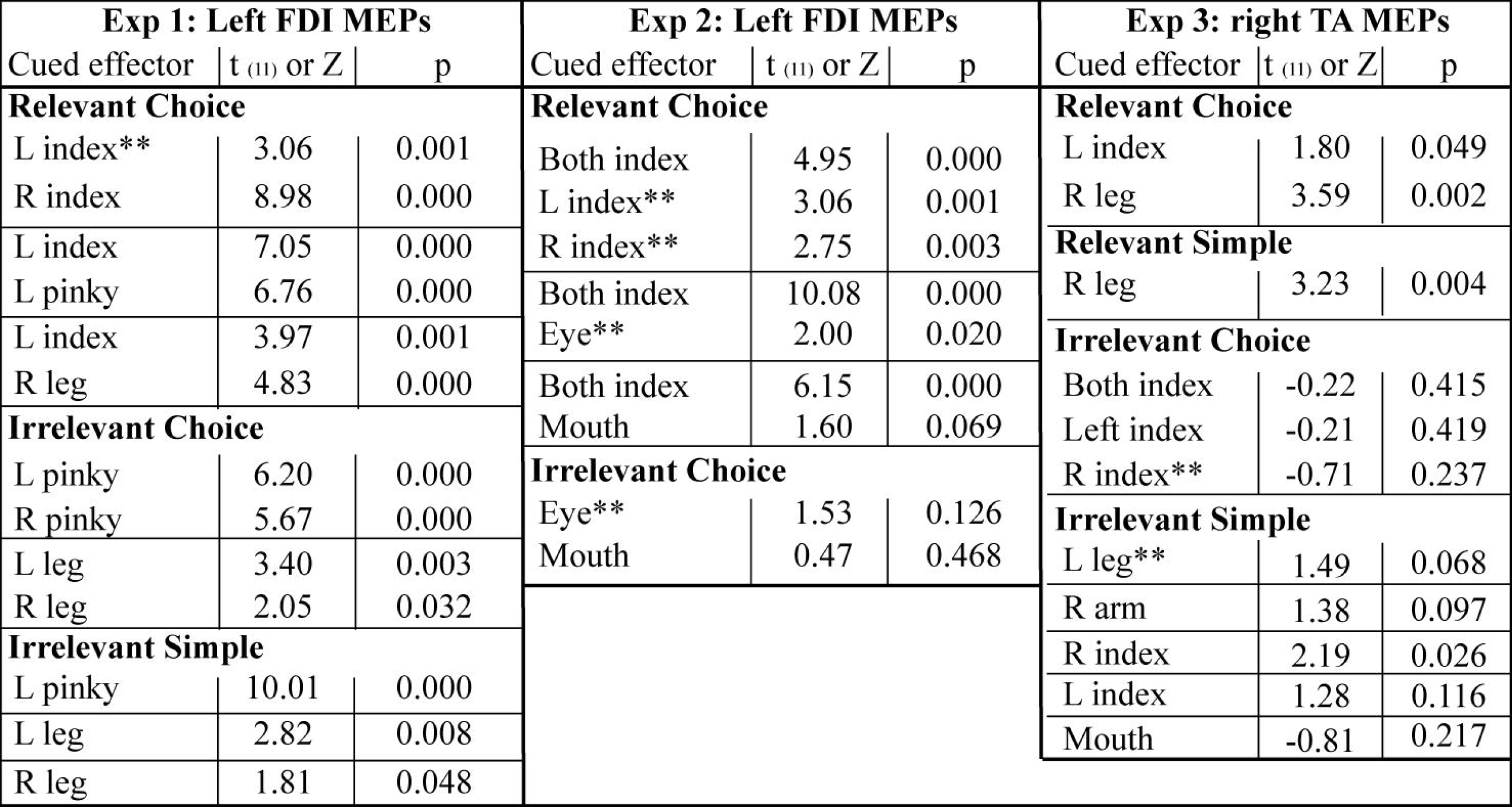
Within-condition test of preparatory inhibition for all three experiments, operationalized as the normalized change in MEP during the delay period relative to the baseline period ((MEP_base_ – MEP_delay_)/MEP_base_). The comparisons were conducted with one-tailed t-tests, motivated by prior studies showing an attenuation of MEPs during the delay period. ** Indicates conditions in which the sample distribution deviated from normality (Shapiro-Wilk test). For these conditions, we present the Z statistic and corresponding p value from the non-parametric Wilcoxon Signed Rank test.

#### Experiment 1

Baseline MEPs for the left FDI averaged 1.02 mV (SD=1.02). Relative to this baseline level, MEPs elicited in the delay period were attenuated in all conditions in which the cue indicated that the participant should prepare a finger movement (all p<0.01, Fig 2). A similar pattern was present when the cue indicated a leg movement (all p<.05). Thus, we observed broad suppression of cortical excitability during response preparation (Greenhouse et al. 2015b), evident when the targeted muscle was part of the task set (Choice Relevant), in most of the conditions in which the muscle was not part of the task set (Choice Irrelevant), and even when there was no choice (Simple Irrelevant RT conditions).

**Figure 2.**
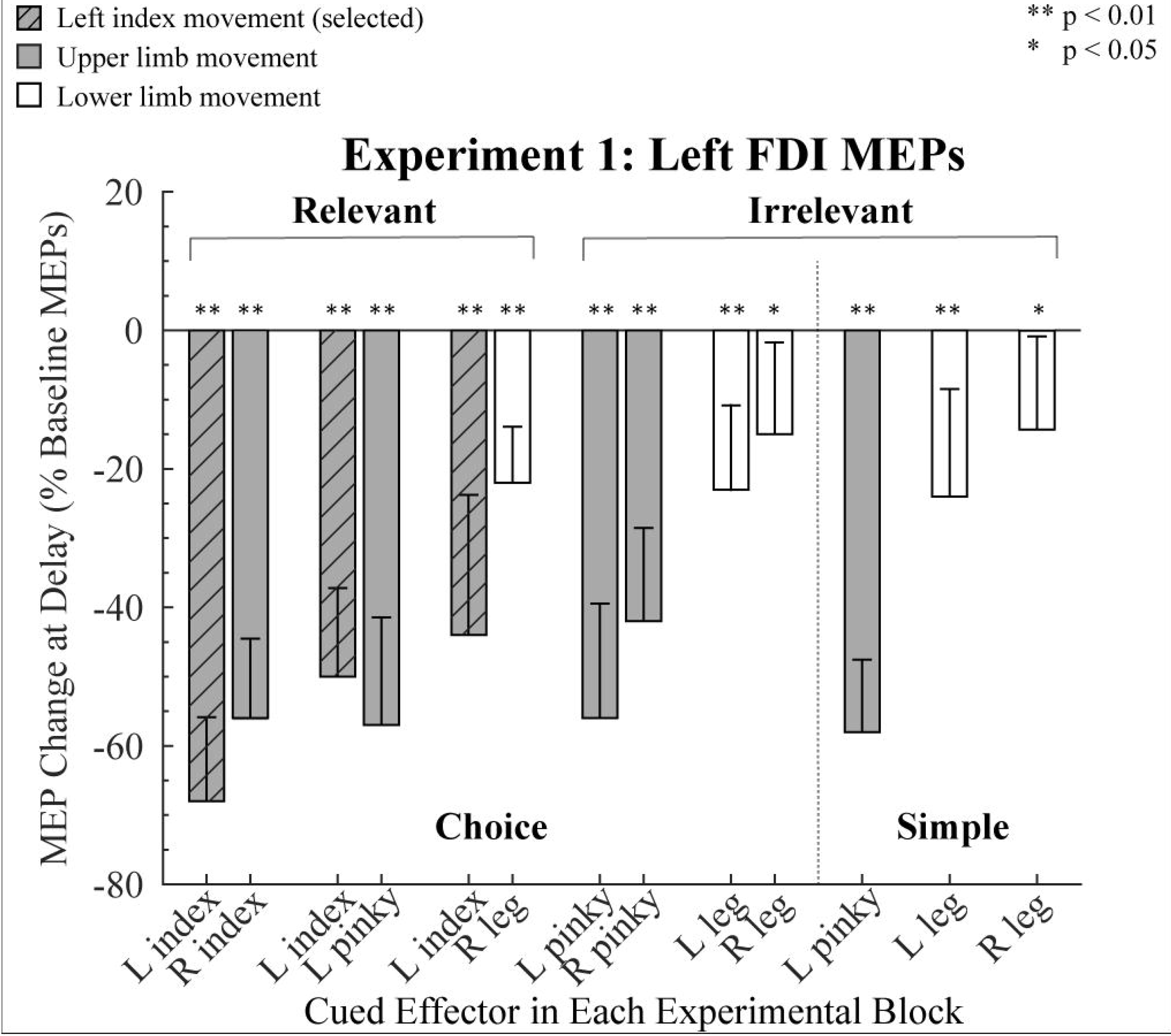
Modulation of MEPs in Experiment 1. MEPs recorded from left FDI during the delay period are expressed as a percentage of baseline (0%). Gray bars indicate trials in which an upper limb movement was cued, and white bars indicate trials in which a lower limb movement was cued. Slashed gray bars indicate when left FDI was the agonist for the forthcoming response. Error bars indicate 95% confidence intervals, depicting if the MEP change during the delay period, relative to baseline, was significantly different than zero (one-tailed test).

To compare the strength of preparatory inhibition between the different experimental conditions, we used a 3 (Task: Choice Relevant, Choice Irrelevant, and Simple Irrelevant) x 2 (Effector: Left Pinky and Right Leg) ANOVA_RM_. We focused on these two effectors since they were included in each of the three types of tasks; the left leg and index fingers were not included in the relevant and irrelevant conditions and, thus, could not be used to test the effect of relevance. The effect of Effector was significant (*F*_(1,11)_ = 42.53, *p* < 0.01, 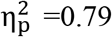), but there was no effect of Task (*F*_(2,22)_ = 3.43, *p* = 0.71, 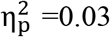), nor an interaction between these factors (*F*_(2,22)_ = 0.40, *p* = 0.67, 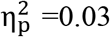). The degree of MEP suppression in left FDI was greater when the cued action required a left pinky movement compared to when it required a right leg movement (mean difference= −39.8% ± 4.4, p< 0.01, Cohen’s d= 6.87). These results indicate that the demands on response selection (Choice vs Simple) and task relevance do not influence the level of preparatory inhibition. However, the magnitude of MEP suppression varied as a function of the movements forming the response set. We recognize that by including the left pinky finger and right leg in the first analysis confounds body segment (upper limb vs lower limb) and body side (left vs right). Given this confound, we performed separate analyses ANOVA_RM_ for each of the tasks (Relevant Choice, Irrelevant Choice, Irrelevant Simple), including in each ANOVA all of the conditions for the task under consideration (see Figure 1).

For the Relevant Choice conditions (Fig 2, left side) we first focused on the three conditions in which the left index finger was cued (selected). The degree of MEP suppression in left FDI varied as a function of the other, non-selected member of the response set (χ^(2)^=8.21, p=0.02). In terms of the post-hoc comparisons, the only reliable difference was that there was stronger suppression of the left FDI when paired with the homologous right index compared to when it was paired with the left pinky (Z=−2.51; p=0.03, r=−0.51). Thus, MEP suppression was greatest in the selected muscle when the choice involved homologous muscles. Second, we examined MEP suppression of left FDI when the left index was not cued (non-selected) in the Choice conditions. Here suppression of left FDI MEPs was weaker when the cued movement was the right leg compared to when the cued movement was either the right index finger (p <0.01, Cohen’s d=1.81) or left pinky (p <0.01, Cohen’s d=1.49). Hence, the amount of left FDI suppression when the left index finger was not selected was stronger when the selected effector was a hand muscle compared to when it was a leg muscle (intra-segment vs inter-segment).

Additional comparisons of anatomy can be made with the data from the Irrelevant conditions in which the left index finger is not part of the response set. For the Choice Irrelevant conditions (Fig 2, middle), we conducted a 2 x 2 ANOVA_RM_ with the factors Body Side (Left, Right) and Effector (Pinky, Foot). There was a main effect for Effector (F_(1,11)_ = 14.88, p<0.01, 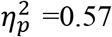), with greater MEP suppression of left FDI when the choice was between two finger movements compared to two leg movements (mean difference = −30±8 %). The effect of Body Side was marginally significant (F(1,11) = 4.82, p = 0.05, 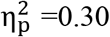), with MEP suppression greater when the forthcoming response was on the left side compared to the right side. The interaction was not significant (F_(1,11)_ = 0.87, p = 0.37, 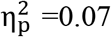). In the Simple Irrelevant conditions (Fig 2, right side), a 1-way ANOVA_RM_ with the factor Competing Effector (Left Pinky, Left Leg, Right Leg) was significant (F_(2,22)_ = 11.44, p<0.01, 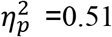). Post-hoc tests showed that left FDI MEP suppression was stronger when participants prepared a left pinky movement compared to a left (p<0.01, Cohen’s d=1.35) or right (p<0.01, Cohen’s d=3.01) leg movement. For the two leg movement conditions, there was no effect of Body Side (p= 0.35, Cohen’s d=1.34).

In summary, the results of Experiment 1 indicate that preparatory inhibition of left FDI is greatest when this muscle is the agonist for the selected response compared to when it is non-selected, replicating earlier results (e.g., Duque and Ivry 2009; Labruna et al. 2014). In terms of our primary question concerning the spread of preparatory inhibition, the magnitude of left FDI MEP suppression was greater when the response set was restricted to finger movements compared to when the response set included a leg muscle. MEP suppression also tended to be greater when the cued response was on the left side of the body compared to when it was on the right side of the body, although this effect was not systematic.

#### Experiment 2

The observations made in Exp 1 are consistent with the hypothesis that the reduced excitability is related to anatomical similarity: MEPs in a hand muscle showed greater suppression when the cued response involved a hand movement compared to when the cued response involved a leg movement. In Exp 2, we further explore anatomical constraints on preparatory inhibition measuring MEPs in left FDI while people prepared finger movements or facial gestures.

Baseline MEPs for the left FDI averaged 0.82 mV (SD=0.51). As in Exp 1, MEPs elicited in the delay period were attenuated in all conditions in which the cue indicated that the participant should prepare a finger movement (all p<0.01, Fig 3). In contrast, when the participants prepared a facial movement, preparatory inhibition in left FDI was only significant in the condition in which the eye movement was prepared in the choice context (Relevant task, p=0.02, see Table 1).

**Figure 3.**
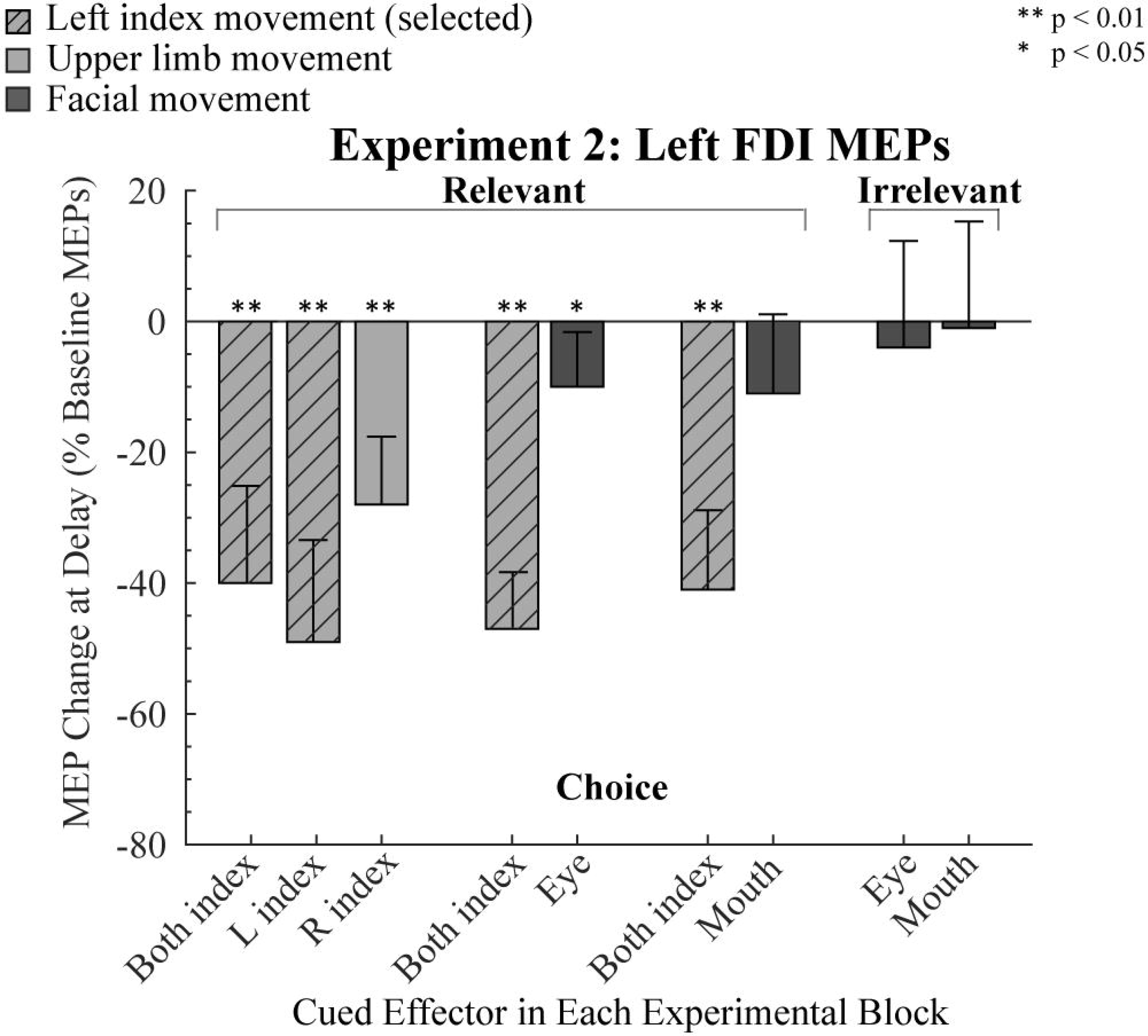
Modulation of MEPs in Experiment 2. MEPs recorded from left FDI during the delay period are expressed as a percentage of baseline (0%). Light gray bars indicate trials in which an upper limb movement was cued and dark gray bars indicate trials in which a facial movement was cued. Slashed gray bars indicate when left FDI was the agonist for the forthcoming response. Error bars indicate 95% confidence intervals, depicting if the MEP change during the delay period, relative to baseline, was significantly different than zero (one-tailed test).

To compare preparatory inhibition between conditions, we first focused on the condition in which the response set was limited to finger movements (Fig 3, left side). Given that the MEP values in a number of conditions violated the normality assumption (see Table 1), the non-parametric Friedman test was used to compare MEP suppression in left FDI when the cued response was for a left index, right index, or bimanual index finger response. There were no significant difference between the three conditions (χ^(2)^ = 5.17, *p* = 0.08,), and planned comparisons showed that the magnitude of MEP suppression in the bimanual condition did not differ from either unimanual condition (left: Z=1.69, p=0.38, r=0.34; right: Z=1.77; p=0.16, r=0.36). The main result to be taken from these analyses is that preparatory inhibition is similar in the bimanual condition compared to the unimanual conditions. We saw this as a prerequisite for the analysis of the facial movement conditions given that the facial gestures are produced bilaterally.

We next compared the three conditions in which participants were cued to prepare a bimanual response (e.g., selected). MEP suppression of left FDI was similar across the conditions (χ ^(2)^ = 0.129, p.>0.94), indicating that the strength of preparatory inhibition was similar when the competing response required a hand or facial movement. However, when the left index finger was not selected, MEP suppression differed across the three conditions (χ ^(2)^ = 6.25, p.>0.04), with the post-hoc comparisons indicating that left FDI was more inhibited when the cue indicated a right index finger movement compared to when the cue indicated an eye movement (Z= 2.51, p=0.03, r-0.51). A similar pattern was observed when the cue indicated a mouth movement, but this comparison did not approach significance (Z=1.84, p=0.18, r=0.38). There was no difference between the mouth and eye movement conditions (Z=0.27; p=2.37, r=0.06). Thus, the results suggest that the suppression of left FDI is reduced when the participants prepared a facial movement. This conclusion is further supported when considering the results from the Irrelevant conditions (figure 3, right). As noted above in the within-condition results, left FDI MEPs in the delay period were not significantly reduced, relative to baseline, when participants had to choose between a mouth or eye movement, and there was no difference between these conditions (Fig 3, right side, Z = −0.55, p = 0.58, r=0.11).

In summary, the results of Exp 2 provide further evidence that the degree of preparatory inhibition varies as a function of the members of the response set. MEP suppression of a hand muscle was greater when the cued response was for a finger movement compared to when the cued response was for a facial movement. In a comparison of the two types of facial responses, we did not observe greater MEP suppression when the participants prepared an eye movement, a strong test of the cortical distance hypothesis. We recognize that the distance from the hand area to the face area may be greater than the extent of preparatory inhibition, an issue we return to in the Discussion. Nonetheless, with this caveat in mind, the results of the first two experiments indicate that the spread of preparatory inhibition is strong within a body segment and weak or absent between segments.

#### Experiment 3

The results of Exps 1 and 2 showed that preparatory inhibition in a finger muscle is much larger when the cued response entails an upper limb movement compared to when the cued response entails a different body segment (lower limb or facial). To ensure that these effects are not specific to upper limb movements, we reversed the situation in Exp 3, measuring MEPs in a leg muscle while participants prepared movements of a leg, finger, or mouth. We opted to stimulate over the left hemisphere, targeting the TA muscle in the right leg. This allowed us to include exact replications of conditions from Exp 1 (Choice: Left Index/Right Leg; Simple: Left Leg), but now with preparatory inhibition probed in a lower limb. As noted above, we only included participants in the main experiment for whom we were able to reliably elicit MEPs in right TA. For these participants, the mean MEPs during baseline were 0.22 mV (SD=0.09), a value that is considerably lower than that for baseline MEPs elicited in FDI in Exps 1 and 2. Nonetheless, we did observe MEPs of at least 0.05 mV on 90% of the trials in the baseline period.

As in the first two experiments, we first conducted within-condition t-tests to assess preparatory inhibition for each condition (Table 1). MEPs elicited in right TA during the delay period were significantly reduced in the two Choice conditions in which the participants prepared a lower limb movement (all p<0.05, Fig 4). A similar trend was observed in the Simple Irrelevant RT condition (p=0.07). In contrast, MEP suppression during the delay period was only observed in two of the seven conditions when an upper limb movement was prepared (left index Relevant Choice and right index Irrelevant Simple, both p<0.05), and was not significant when a mouth response was prepared. Thus, preparatory inhibition in right TA was robust when participants prepared a leg movement (right or left leg), but inconsistent or absent when preparing an upper limb movement or facial gesture.

**Figure 4.**
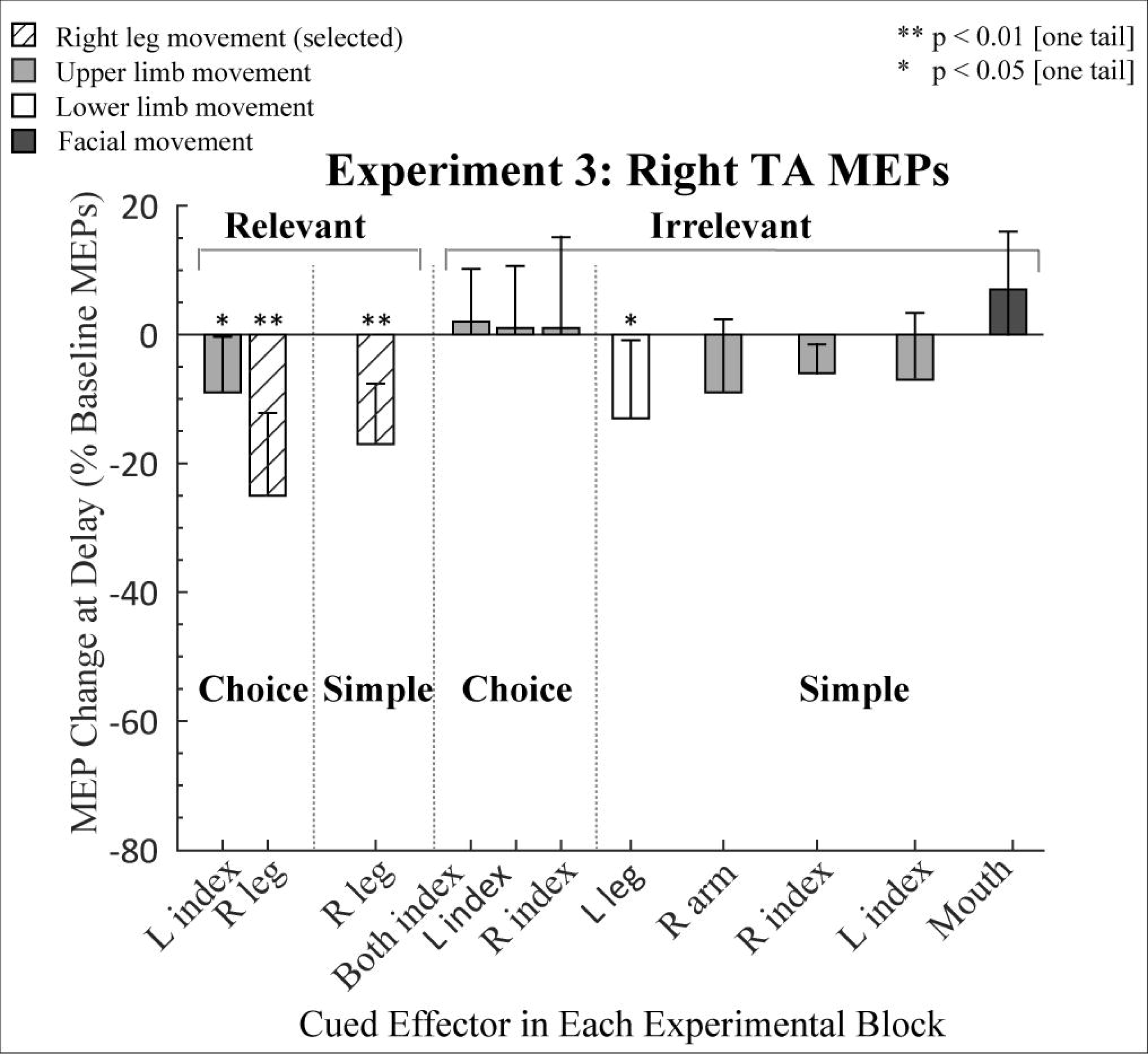
Modulation of MEPs in Experiment 3. MEPs recorded from right TA during the delay period are expressed as a percentage of baseline (0%). Light gray, white, and dark gray bars indicate trials in which the cued response required an upper limb, lower limb, or facial movement, respectively. Slashed white bars indicate when right TA was the agonist for the forthcoming response. Error bars indicate 95% confidence intervals, depicting if the MEP change during the delay period, relative to baseline, was significantly different than zero (one-tailed test).

Turning to the between-condition comparisons, preparatory inhibition in right TA was greater when that muscle was selected for the forthcoming response compared to when it was not selected (Choice Relevant: t_(12)_ = 3.14, p = 0.01, Cohen’s d=1.19). No differences were found when the right leg was selected as part of either a Choice or a Simple task (t_(12)_ = −1.04, p = 0.32 Cohen’s d=0.35), consistent with the results of the first experiment, indicating that preparatory inhibition is independent of the task context.

For the Irrelevant conditions, we conducted three analyses to compare preparatory inhibition in the right TA when the cued response a different lower limb effector to conditions in which the cued response was from another body segment. For the former, we used left leg movements; for the latter, the cued response either involved upper limb effectors or the mouth. First, we compared the left leg condition to the upper limb condition, taking the average of the three upper limb effectors in the Choice condition. This contrast was significant (Z = −2.20, p = 0.03, r=0.45,), with greater MEP suppression in right TA when the selected limb was from the same body segment. The second contrast was between the left leg and the average of the three upper limb effectors in the Simple conditions. Here the difference was not significant (Z = −0.39, p = 0.7, r=0.08,). The third contrast, between the left leg and mouth approached significance (Z = − 1.82, p = 0.07, r=0.37,).

Overall, the results of Exp 3 are consistent with the idea that anatomical constraints on preparatory inhibition are not specific to upper limb muscles, but also hold for lower limb muscles. This prediction was supported by two of the contrasts of different body segments; it was not supported by the third (lower vs. upper segment, Simple conditions). We note that our sensitivity in this experiment is reduced given the relatively low MEPs elicited from right TA.

### Reaction Times

RTs were relatively fast (around 250 ms), indicating that the participants had used the cues to prepare the forthcoming response during the delay period (Fig 5). This is most clearly evident in the comparison of Choice and Simple RTs for each effector in Exps 1 and 3: Mean RTs in the Choice RT conditions were similar to those observed in the Simple RT conditions. The difference scores ranged from 0 ms to 22 ms, and even the largest difference (Exp 3, right index finger) was not significant (p=0.35). RTs were also similar on trials in which the TMS pulse was applied just prior to the start of the trial (baseline) or when applied during the delay period in all three experiments (p > 0.10), with data collapsed across conditions.

**Figure 5.**
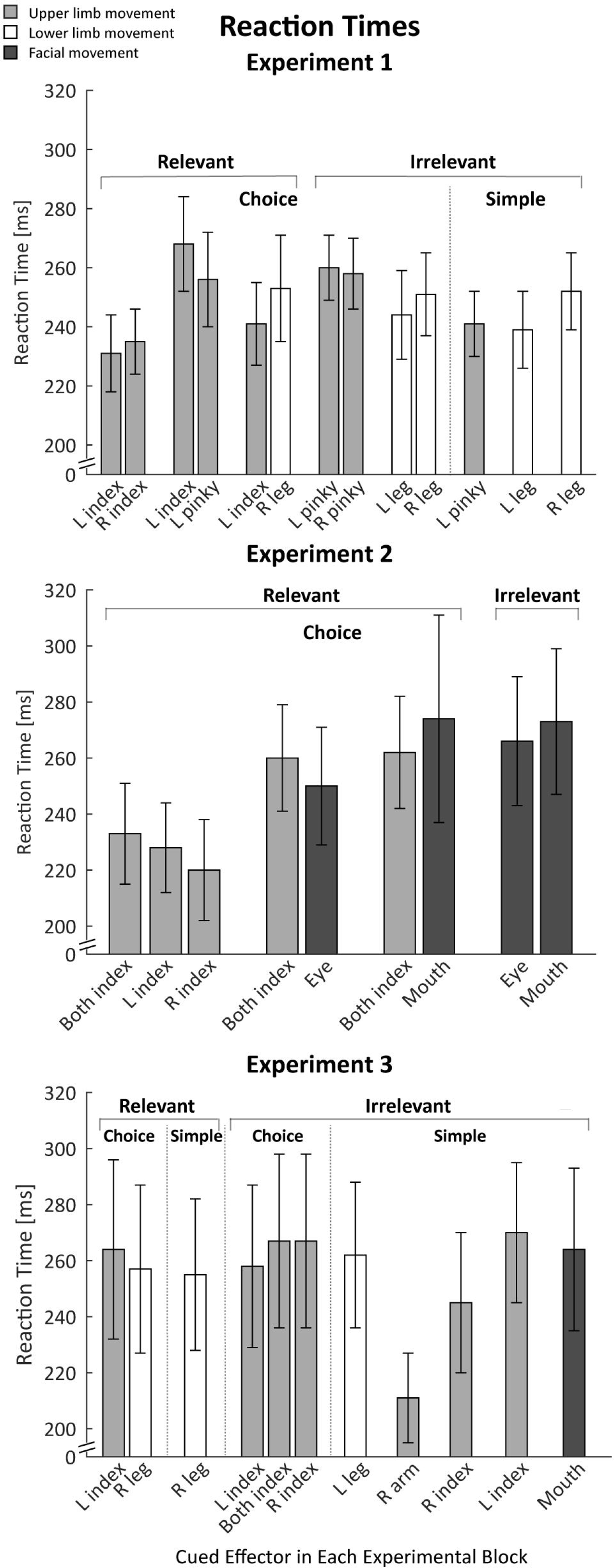
Reaction times for Experiments 1-3, combining trials in which TMS was applied at baseline and during the delay period. Light gray, white, and dark gray bars indicate trials in which the cued response required an upper limb, lower limb, or facial movement, respectively. Error bars indicate SEMs.

There were some effector-specific effects on RT. For example, we can compare left and right sided RTs for the index finger, pinky, and leg in three Choice conditions in Exp1 (Fig 5, top). Mean RTs were fastest for index finger movements (233±12 ms), followed by leg movements (247±15 ms), and slowest for pinky movements (259±10 ms). However, a 3 (Effector) x 2 (Side) ANOVA_RM_ showed that these differences were not significant (all p>0.14).

In the Choice RT conditions, the RT for a given effector was modulated by the other member of the response set. For example, a 1-way ANOVA_RM_ on the RTs for the left index finger in the three Choice conditions showed a main effect (F_(2,22)_ = 7.60, p < 0.01, 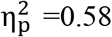), with slower RTs when the left index finger movement was paired with the pinky of the same hand, compared to when it was paired with the right Index finger (p=0.01, Cohen’s d=2.53) or with the right Leg (p=0.03, Cohen’s d=1.82). This pattern suggests that the participants adopted, to some degree, a task set in which the speed of movement initiation for a given condition was relatively constant for each choice, adjusted to the rate of the slower member of the response pair.

A similar pattern was evident in Exp 2 (Fig 5, middle). RTs were slower for the facial gestures compared to the finger responses. Focusing on the 3-choice condition that involved bimanual responses (averaging RTs over left and right fingers since the responses were tightly coupled), finger RTs were slower in blocks in which these responses were paired with facial responses than with a unimanual finger response (mean difference with eye 27±49 and with mouth 29±48), although the ANOVA_RM_ showed only a marginal effect (p= 0.07, 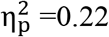).

In Exp 3 (Fig 5, bottom), finger RTs in the Choice conditions were relatively invariant, with no advantage in conditions in which all responses were with the fingers compared to when a finger and leg response were paired. At first glance, RTs in Exp 3 were slower than in the first two experiments. In a post-hoc analysis, we compared RTs for the left index finger across experiments, focusing on this finger since it was the only effector paired in all three experiments with another upper limb effector. The outcome of this 1-way ANOVA was not significant (p=0.52, 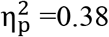).

RT was not related to the magnitude of the MEPs (see Duque et al. 2017 for a discussion on this issue), similar to what has been observed in previous studies (but see Hannah et al. 2018). This can be seen in a comparison between conditions: For instance, in Exp 2, bimanual RTs tended to be slower when paired with facial responses than when paired with unimanual finger responses, but MEPs elicited from the left FDI were relatively invariant across conditions. Even more compelling, it is not observed in a trial-by-trial analysis performed on an individual basis. Pooling across conditions involving the left index finger, there was no consistent pattern of correlation between RT and MEP for the left index finger.

## Discussion

Preparing to move entails the recruitment of inhibitory mechanisms. This preparatory inhibition is evidenced by the attenuation of MEPs elicited during a delay period when participants prepare to initiate a cued response. Several studies have identified constraints on the magnitude of this phenomenon; for example, the degree of MEP suppression is modulated by task difficulty (Beck and Hallett 2010; Greenhouse et al. 2015a; Klein et al. 2014). These findings indicate that preparatory inhibition is not generic. In the current study, we extend this work, systematically examining anatomical constraints on preparatory inhibition.

### Anatomical Constraints on Preparatory Inhibition

Consistent with previous findings, preparatory inhibition was generally greatest when the targeted muscle was the agonist for the forthcoming movement. Moreover, the magnitude of MEP suppression for the selected conditions was independent of the other member of the response set. This was most evident in Exp 2 where MEP suppression in the left FDI was similar across Choice conditions in which the left index finger was paired with the right index finger or paired with an eye or mouth movement.

A different pattern was observed when the cue indicated a response other than the left index finger. Preparatory inhibition in left FDI was pronounced if that effector was from the same body segment (e.g., another manual response), but much weaker if the cued effector was from a different body segment. In Exp 1, the mean level of MEP suppression, relative to baseline was − 42% when the cued response involved another finger movement and only −20% when the cued response involved a leg movement. Similarly, in Exp 2, MEPs were reduced by −28% when the cue indicated a right index finger movement and only reduced by −8% when the cue indicated a facial movement. Indeed, in the latter experiment, mean MEP amplitudes were not significantly different from baseline in three of the conditions involving facial responses.

This pattern was similar for conditions in which the left index finger was relevant or irrelevant. Moreover, the magnitude of preparatory inhibition did not depend on whether the cue required a decision between alternative responses (Choice Conditions) or always specified the same response (Simple Conditions). For example, on trials in Exp 1 in which the planned response was with the left pinky finger, MEP suppression of left FDI was similar when the left index finger was part of the response set or not part of the response set. Consistent with the results reported in Greenhouse (2015), the magnitude of preparatory inhibition does not appear to depend on task relevance or choice behavior.

Taken together, the results of Exps 1 and 2 indicate that the magnitude of preparatory inhibition targeted at non-responding effectors is greater when the planned response is from the same body segment (e.g., hand) compared to when it entails a different body segment (leg or face). To test the generality of this hypothesis, the TMS probe was directed at right TA, the agonist for adduction movements of the lower leg, in Exp 3. Here we also included conditions in which the response set either included or didn’t include the right leg. The pattern was similar to that observed in Exps 1 and 2. MEPs from right TA were significantly suppressed during the delay period when the cue called for the preparation of either a right or left leg movement. In contrast, MEP suppression of right TA was reduced or absent when the cue indicated a hand, arm, or facial movement.

Qualitatively the magnitude of preparatory inhibition appears to be lower for right TA compared to left FDI. We are hesitant to draw any inferences concerning this pattern. First, this between-experiment comparison confounds side and segment, given our decision to focus on right TA. Second, although we normalize our measure of preparatory inhibition by expressing the change in the delay period relative to baseline, it is important to keep in mind that MEPs are much more difficult to obtain from leg muscles, and when obtained, are weaker than those elicited from FDI (Kesar et al. 2018). Most important, the claims about intra- vs intersegment differences are evident in the within-experiment comparisons where the TMS probes are always restricted to the same muscle.

### Anatomy vs. Function

We interpret the current results to indicate that the extent of preparatory inhibition is constrained by anatomy, dropping in strength when the distance between the selected effector and the muscle targeted by TMS is increased. One variant of this distance hypothesis is that the extent of preparatory inhibition may be related to the motor homunculus. Exp 2 was designed to test this hypothesis, building on the fact that the hand area is closer to the cortical representation of the eyes compared to the cortical representation of the mouth. The results of experiment 2 failed to support this strong version of the cortical distance hypothesis: When either a squint or smile were planned, there was minimal change in left FDI MEPs, and numerically, the small effects were comparable for the two types of facial gestures.

However, there are a number of caveats to keep in mind when considering the cortical distance hypothesis. First is the general concern with all null results. Second, although the eye representation is closer to the hand area, the distance is still relatively large, at least in comparison to the distance between finger representations (Meier et al. 2008; Weiss et al. 2013). It may be that the spread of excitability changes does follow a cortical gradient, but that it is negligible beyond some maximal distance. A finer-grained analysis would be required to test the cortical distance hypothesis; for example, compare the magnitude of preparatory inhibition in left FDI in conditions in which the cue specifies a finger, wrist, lower arm, or upper arm movement.

The current results do reveal a consistent difference between conditions in which the planned movement is from the same body segment (lower, upper, face) or a different body segment, with the former producing greater reduced excitability in the probed muscle. Rather than attribute these effects to the cortical distance of motor representations, the difference may reflect the synergistic recruitment of intrasegmental representations. The motor homunculus visualized across the motor cortex is recognized as a simplification given that there is considerable overlap between motor representations. Indeed, it has been proposed that a clear spatial separation is limited to representations of different body segments (Schieber 2001; Zeharia et al. 2012). By this view, the interactions within a segment in terms of preparatory inhibition could arise from the fact that the fingers of one hand, or even fingers between two hands, are frequently co-activated for a given movement. Preparatory inhibition might extend to effectors within the same body segment as the cued one to reduce activation of muscles that are close in cortical space to the agonist for the forthcoming movement.

There are well-defined movements that do involve intersegmental coordination. For example, when reaching for objects, the eyes and hands move in a coordinated manner, and some of the ethological gestures described by Graziano and colleagues (Desmurget et al. 2014; Fernandino and Iacoboni 2010; Graziano 2016) involve coordinated movements between the upper limbs and face (e.g., eating). The fact that our results failed to reveal consistent MEP suppression in FDI when preparing facial gestures argues against these function-based hypotheses. Similarly, in a preliminary study (Labruna et al. 2016), we tested experienced drummers to see if they showed greater preparatory inhibition in an upper limb when preparing a leg movement given that drumming requires extensive intersegmental coordination. The data from this group was similar to that reported here, with minimal MEP suppression of FDI when the drummers prepared a leg movement. In summary, the present picture suggests that the spread of preparatory inhibition is best defined in terms of a segmental criterion, rather than one based on functional considerations.

### Implications for Models of Preparatory Inhibition

A recent review by Duque at al. (2017) summarizes three functional models of preparatory inhibition. The first of these models suggests that inhibition is restricted to task-relevant muscles, reflecting a competition between candidate effectors (Duque et al. 2005, 2010). The second model suggests preparatory inhibition arises from the operation of two processes, one producing a global or broad inhibitory effect and the other focused at only the selected response representation. The third model emphasizes a single process that operates in the form of an ‘spotlight’ centered over the selected response representation with the width of the aperture constrained by the task context such as whether or not selection entails a choice (Greenhouse et al. 2015b). According to the spotlight model, inhibition, or reduced excitation, facilitates the selection and initiation of motor responses by reducing background noise and, thus, increasing the gain within the motor system.

We observed preparatory inhibition, independent of whether the probed muscle was part of the response set or was task irrelevant. Moreover, we also observed robust MEP suppression when the probed muscle was the sole member of the response set. These findings are at odds with the competition model since competition is absent in the task-irrelevant conditions and Simple conditions. In contrast, the two-process and spotlight models are consistent with the current findings, although we suggest an additional anatomical constraint on preparatory inhibition. A spotlight might operate at the level of body segments, with the strongest influence over the body segment that includes the selected response representation, and negligible effect on representations from other body segments. With respect to the two-process model, the current results would indicate that the process producing a broad reduction of excitability is not generic. Rather, its extent appears to be categorical and mostly limited to muscles within the same body segment as the agonist effector. The notion of a categorical constraint based on body segment, however, should be qualified given that we may lack the sensitivity to detect effects in the tail of a gradient, one that spans large cortical distances.

In terms of function, the current data do not differentiate hypotheses that focus on how preparatory inhibition might prevent premature responses or facilitate gain modulation during response planning. Future work may be able to capitalize on the spatial constraints identified here to better address functional questions.

### Relationship of anatomical constraints in preparatory and reactive inhibition

TMS has been used to characterize the dynamics of cortical excitability in tasks involving reactive inhibition, such as the stop-signal task in which a planned response is aborted. One prominent idea is that, when the stop signal requires the termination of all volitional movement (where the planned response involves one or more effectors), the inhibitory signal is broadcast in a global manner, manifest in both task relevant and task-irrelevant muscles (Badry et al. 2009; Coxon et al. 2006; Greenhouse et al. 2012; Leocani et al. 2000; Majid et al. 2012). Most relevant to the present discussion, reactive inhibition is seen in both intra- and intersegmental muscles.

Superficially, it may appear that preparatory and reactive inhibition arise from different processes given that we find, at best, modest preparatory inhibition between body segments whereas reactive inhibition tasks point to a global process. However, it remains unclear if the TMS data provide strong evidence of a difference between preparatory and reactive inhibition. Similar to the effects observed here, the magnitude of reactive inhibition in task-irrelevant muscles is much larger for intrasegmental muscles compared to intersegmental muscles. For example, Badry et al. (2009) used TMS to elicit MEPs in either the thumb or leg after a stop signal had indicated that the participants should abort an index finger response. Relative to baseline, thumb MEPs were reduced by close to 50%, whereas leg MEPs were only reduced by 15% (see also, Greenhouse et al. 2012; Majid et al. 2012). Similarly, stopping speech resulted in only a 15% reduction in hand MEPs (Cai et al. 2012).

In sum, the stop signal literature also points to a gradient in the extent of reactive inhibition, similar to that observed here with preparatory inhibition, with only weak changes in corticospinal excitability when the probed muscle is at a different segmental level as the task relevant effector. This observation by itself offers only weak evidence for a common mechanism underlying preparatory and reactive stopping. Future studies can be designed to provide more direct tests. Whereas studies using a range of methods have detailed a cortico-basal ganglia circuit recruited for reactive stopping, similar work is needed to understand the networks that result in preparatory inhibition.

### Conclusions

The three experiments reported here provide converging evidence that preparatory inhibition is constrained by anatomy. A marked reduction in corticospinal excitability was observed when the response involved a muscle from the same body segment, and reduced or even absent when the response involved a muscle from a different body segment. These results are consistent with models in which an inhibitory process is targeted at specific motor representations, with a spatial extent limited to motor representations within the same body segment.

## Acknowledgments

This work was supported by grants from the Belgian National Funds for Scientific Research (FRS-FNRS: MIS F.4512.14), the Fondation Médicale Reine Elisabeth (FMRE), the National Institute of Health (NS092079, NS097480), and the France-Berkeley fund. We thank Simone Ewell-Szabo for drawing Figure 1a.

